# Reading instruction causes changes in category-selective visual cortex

**DOI:** 10.1101/2022.02.02.477919

**Authors:** Jason D. Yeatman, Daniel R. McCloy, Sendy Caffarra, Maggie D. Clarke, Suzanne Ender, Liesbeth Gijbels, Sung Jun Joo, Emily C. Kubota, Patricia K. Kuhl, Eric Larson, Gabrielle O’Brien, Erica R. Peterson, Megumi E. Takada, Samu Taulu

## Abstract

Education sculpts specialized neural circuits for skills like reading that are critical to success in modern society but were not anticipated by the selective pressures of evolution. Does the emergence of brain regions that selectively process novel visual stimuli like words occur at the expense of cortical representations of other stimuli like faces and objects? To answer this question we conducted a randomized controlled trial with pre-school children (five years of age). We found that being taught reading versus oral language skills induced different patterns of change in category-selective regions of visual cortex. Reading instruction enhanced the response to text but did not diminish the response to other categories. How these changes play out over a longer timescale is still unknown but, based on these data, we can surmise that high-level visual cortex undergoes rapid changes as children enter school and begin establishing new skills like literacy.

## Introduction

Through education, humans are able to master new skills, such as literacy, that were not anticipated by the selective pressures of evolution. How is the human brain able to automate an evolutionarily novel function like reading such that it becomes as effortless and automatic as evolutionarily older functions such as face recognition? Literacy is the paradigmatic example of how acquiring expertise in a new domain prompts the development of specialized neural circuits that are at least partially dedicated to the computations required for the new skill (Dehaene et al., 2015; Yeatman and White, 2021).

The “neuronal recycling hypothesis” is the dominant framework for explaining the emergence of cortical regions that are specialized for evolutionarily novel, uniquely human cultural inventions such as literacy (Dehaene and Cohen, 2007). This framework posits that over the course of learning, the computations required for the novel function find their neuronal niche in a related circuit (high level visual cortex in the case of literacy), and that the learning process involves competition between the novel function (e.g., word recognition) and an evolutionarily older function (e.g., face or object recognition). The core tenet of neuronal recycling as it applies to literacy is competition between words and faces (or objects and bodies) for cortical territory in ventral occipito-temporal cortex (VOTC) such that regions of cortex that would otherwise be specialized for processing faces or objects are, instead, tuned to words. This theoretical framework is supported by the observation that, within the mosaic of regions in high level visual cortex that selectively process specific visual categories such as faces, bodies, objects and scenes, there exists a region that selectively processes visual words (Cohen et al., 2002; Grill-Spector and Weiner, 2014; Malach et al., 2002). This “visual word form area” (VWFA) is localized to the same region of VOTC in literate people across languages, cultures and orthographies (Dehaene et al., 2015; Dehaene and Cohen, 2007; Nakamura et al., 2012; Rueckl et al., 2015), its response properties are correlated with reading skills (Ben-Shachar et al., 2011; Kubota et al., 2019), and its characteristic response to text stimuli occurs in literate but not illiterate adults (Dehaene et al., 2015, 2011; Hervais-Adelman et al., 2019).

There is widespread correlational evidence supporting the neuronal recycling hypothesis. For example, comparisons between literate and illiterate adults suggest that the patch of cortex occupied by the VWFA in literates responds to faces in illiterates (Dehaene et al., 2011), and over the typical course of schooling the VWFA becomes increasingly selective for words compared to other visual categories (Cantlon et al., 2010; Kubota et al., 2019). The left hemisphere specialization for the emergence of literacy has even been offered as an explanation for the right hemisphere specialization for face recognition (Behrmann and Plaut, 2020). On the other hand, recent observations suggest that the VWFA emerges in a patch of cortex that is not otherwise tuned for any specific stimulus class (Dehaene-Lambertz et al., 2018; Hervais-Adelman et al., 2019; Saygin et al., 2016). Moreover, literacy is associated with improved face and object recognition performance challenging the notion of competition between visual categories (van Paridon et al., 2021). These data have prompted a reexamination of the neuronal recycling hypothesis by suggesting that word-selective responses do not compete for cortical territory with other visual categories. However, a recent longitudinal study revealed that limb-selective regions in VOTC lose cortical territory to word-selective regions over the course of elementary school (Nordt et al., 2021).

Given the myriad changes that a child (or adult) undergoes when they enter school, isolating the causal effects of literacy learning is impossible without a randomized controlled trial (RCT) design. Despite the hundreds of correlational studies examining the neural underpinnings of literacy, a causal test of neuronal recycling is lacking.

Causal evidence of neuronal recycling requires the demonstration that literacy learning involves competition between words and other visual categories for cortical territory; the critical test is an RCT where pre-school children are randomly assigned to either a literacy intervention or a control intervention. Here we ran such an RCT focused on the initial phase of literacy learning where children begin learning to recognize letters, associate letters with sounds, and blend sounds to decode simple words. Pre-literate children (4y10m to 6y2m) were randomly assigned to an intervention program involving direct instruction in either (a) letters, sounds and the foundations of literacy (Letter Intervention); or (b) oral language comprehension, grammar and vocabulary (Language Intervention). Both programs were delivered by experienced teachers in small groups (6 children; 2 teachers) 3 hours a day, five days a week for two weeks (30 hours total) during the summer before kindergarten. Both intervention programs followed an identical schedule that involved: (1) direct instruction in new concepts (either letters or oral language); (2) teacher monitored practice through targeted activities that provide multisensory experience with the new concept; (3) guided repetition and practice; (4) play breaks to maintain focus and promote learning. The Letter Intervention curriculum included systematic instruction in the form and sound of each letter, practice creating and identifying the letter in multiple modalities (e.g., pen and paper, clay, in books), blending letters to form words and phonological awareness. The Language Intervention (control) systematically taught vocabulary, oral language comprehension and grammar skills, maintaining a matched lesson plan in terms of the balance between direct instruction, individual practice and play, but without any exposure to text.

Magnetoencephalography (MEG) and source localization (based on individual anatomical MRI data) were used to measure cortical responses to words, faces and objects (**Figure 1**) immediately before and after the intervention program. Neuronal recycling predicts that: (1) the two intervention programs will induce different changes in selectivity within the patch of cortex that is destined to become the VWFA (threeway intervention group by stimulus condition by time point interaction), and (2) the Letter Intervention group will show an increased response to words at the expense of faces or objects. We address the first question based on data from the RCT and then, to address the second question with greater statistical power, we combine data from a second, independent cohort that participated in the Letter Intervention.

**Figure 1:**
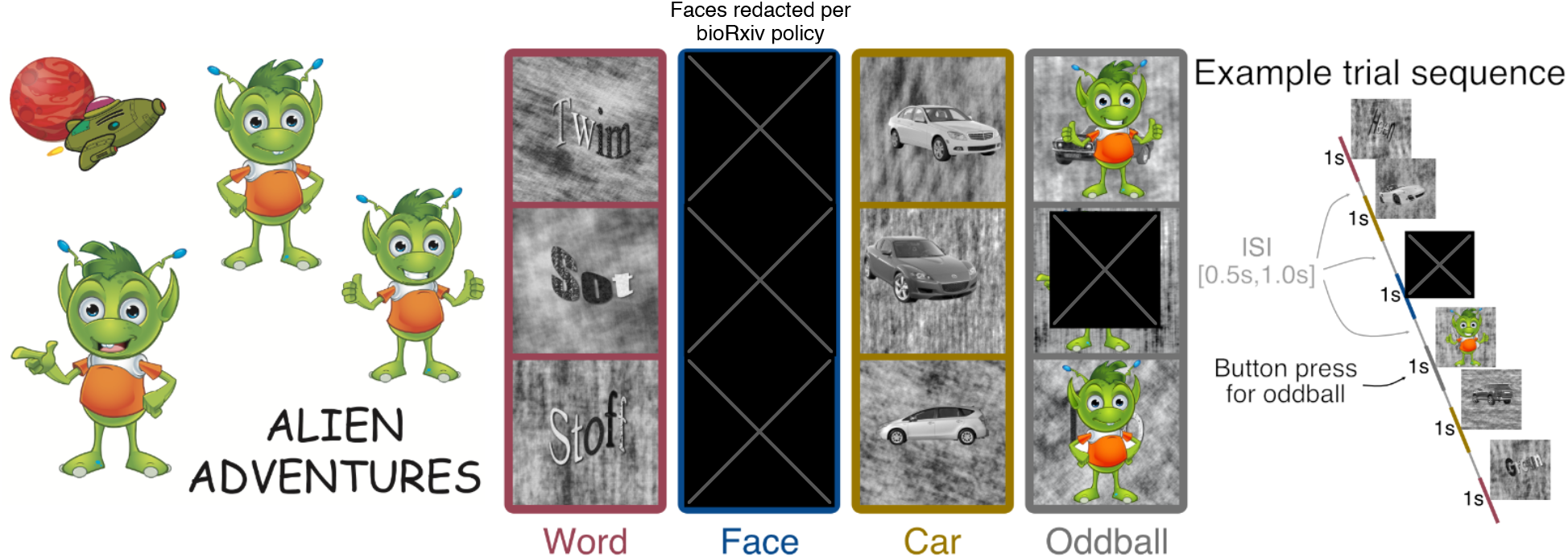
Stimuli and procedure for MEG experiment. Children played the game “Alien Adventure” in which they searched for a friendly alien, hidden in a sequence of images. Participants were instructed to focus on the center of the screen and press a button each time the alien (oddball) appears. Images of pronounceable pseudowords rendered in an unfamiliar font (hereafter “Words”), children’s faces, and cars (30 each) were displayed in a random order with ten oddball trials occurring at random times during the experiment. Oddball trials were excluded from the analysis. Faces were selected as a comparison condition since face-selective regions are immediately adjacent to word-selective regions (in literate brains) and competition between words and faces is the foundation of the original instantiation of the neuronal recycling hypothesis (Behrmann and Plaut, 2020; Dehaene et al., 2011; Dehaene and Cohen, 2007). Cars were chosen as the second comparison condition to test the notion that competition occurs more generally for object-selective cortex (with empirical data demonstrating that cars serve as a good baseline comparison for words (Kubota et al., 2019); see Methods for more details).

## Results

### Two weeks of literacy training improves alphabet knowledge and decoding skills

To test the efficacy of the intervention programs we compared pre-test and post-test scores for the Letter versus Language intervention groups using a linear mixed effects (LME) model with fixed effects of time (post vs. pre), fixed effects of intervention group (Letter vs. Language), a group by time interaction, and random intercepts and slopes for each participant (Barr et al., 2013; Bates et al., 2007). Comparing the change in behavioral measures between the intervention groups revealed significant group by time interactions for the two primary outcome measures (**Figure 2**): lower case Letter Knowledge (the number of letters and sounds a child could name; t(92) = 2.97, p = 0.004) and Decoding (the number of non-word three-letter sequences a child could pronounce; t(92) = 3.07, p = 0.002). Moreover, the Letter intervention generalized to growth in upper case Letter Knowledge even though the training focused on lower case letters (group by time interaction, t(92) = 2.68, p = 0.008). Modeling intervention-driven growth separately for each group confirmed that the improvements were specific to the Letter Intervention group: Over the two-week Letter Intervention, children made significant gains in Letter Knowledge (t(46) = 4.49, p = 0.000046, **Figure 2**) and Decoding (t(46) = 3.29, p = 0.0019). Children in the Language intervention did not show changes in Letter Knowledge (t(46) = 0.76, p = 0.44) or Decoding (t(46) = -0.83, p = 0.41).

**Figure 2:**
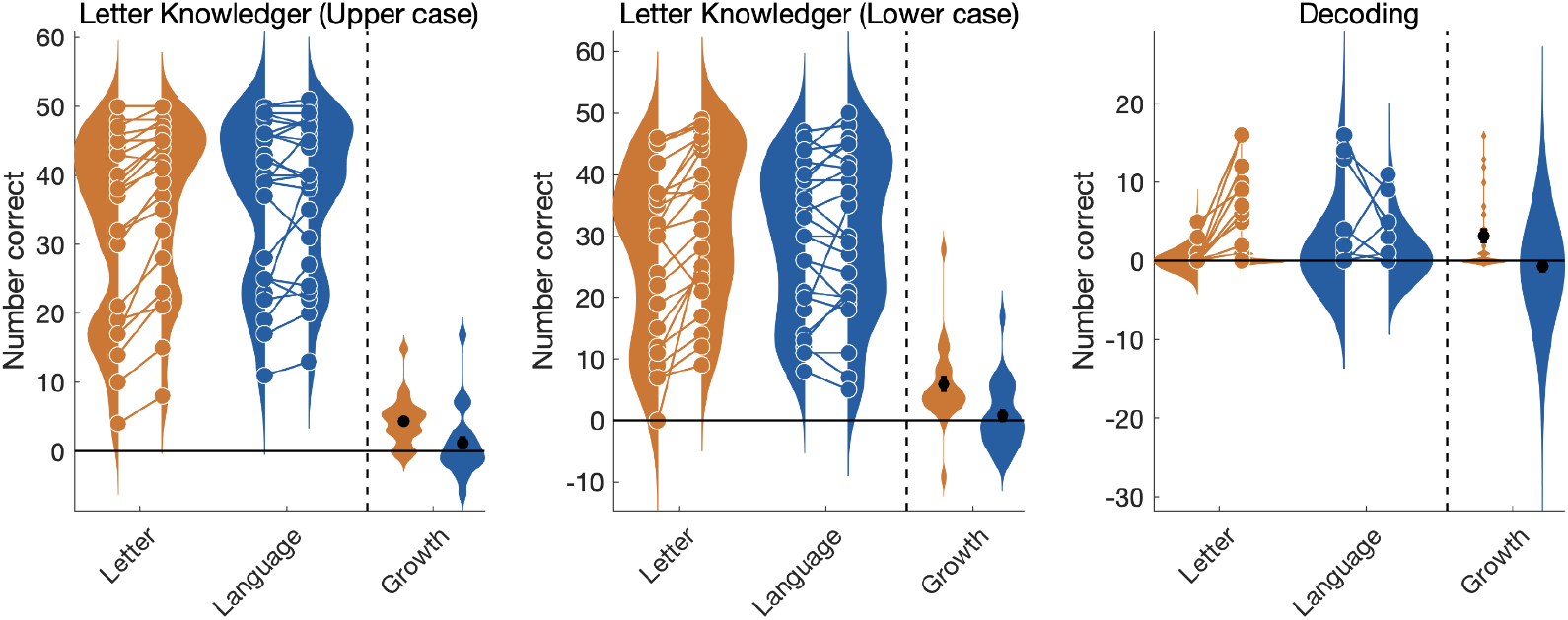
Improvement in Letter Knowledge and Decoding is specific to Letter Intervention. Violin plots show individual scores pre and post-intervention for the Letter (orange) and Language (blue) groups overlaid on a smoothed kernel density plot. For each measure, growth is calculated as the difference between each participant’s post versus pre intervention score. Distributions of growth are shown with a black bar representing the mean growth (± 1 standard error) for each group.

### Intervention-driven changes in visual cortex

To test the hypothesis that two weeks of literacy training changes the tuning of VOTC to words, faces, and objects, we fit a LME model to source reconstructed MEG responses within an *a priori* anatomically defined region of interest (ROI) encompassing the typical location of the VWFA (**Figure 3A**). The ROI (hereafter referred to as word-selective cortex) was taken from a probabilistic functional atlas of word-selective regions in visual cortex (Rosenke et al., 2021). Focusing on a region that has already been established as word-selective in literate adults allows us to investigate the emergence of its function during the early stages of learning. To limit the number of statistical comparisons, we computed the grand average waveform by averaging MEG responses localized to that ROI across stimulus conditions and subjects, identified the first peak (which occurred at 175ms), and defined a time window of ± 50ms around the peak. We then averaged responses within this 135ms-235ms time window within subjects separately for each condition (Words, Faces, Cars), and compared responses across conditions and intervention groups. This early time window corresponds to the stimulus-evoked visual response in VOTC (Hirshorn et al., 2016; Marinkovic et al., 2003; Thesen et al., 2012; Woolnough et al., 2020).

**Figure 3:**
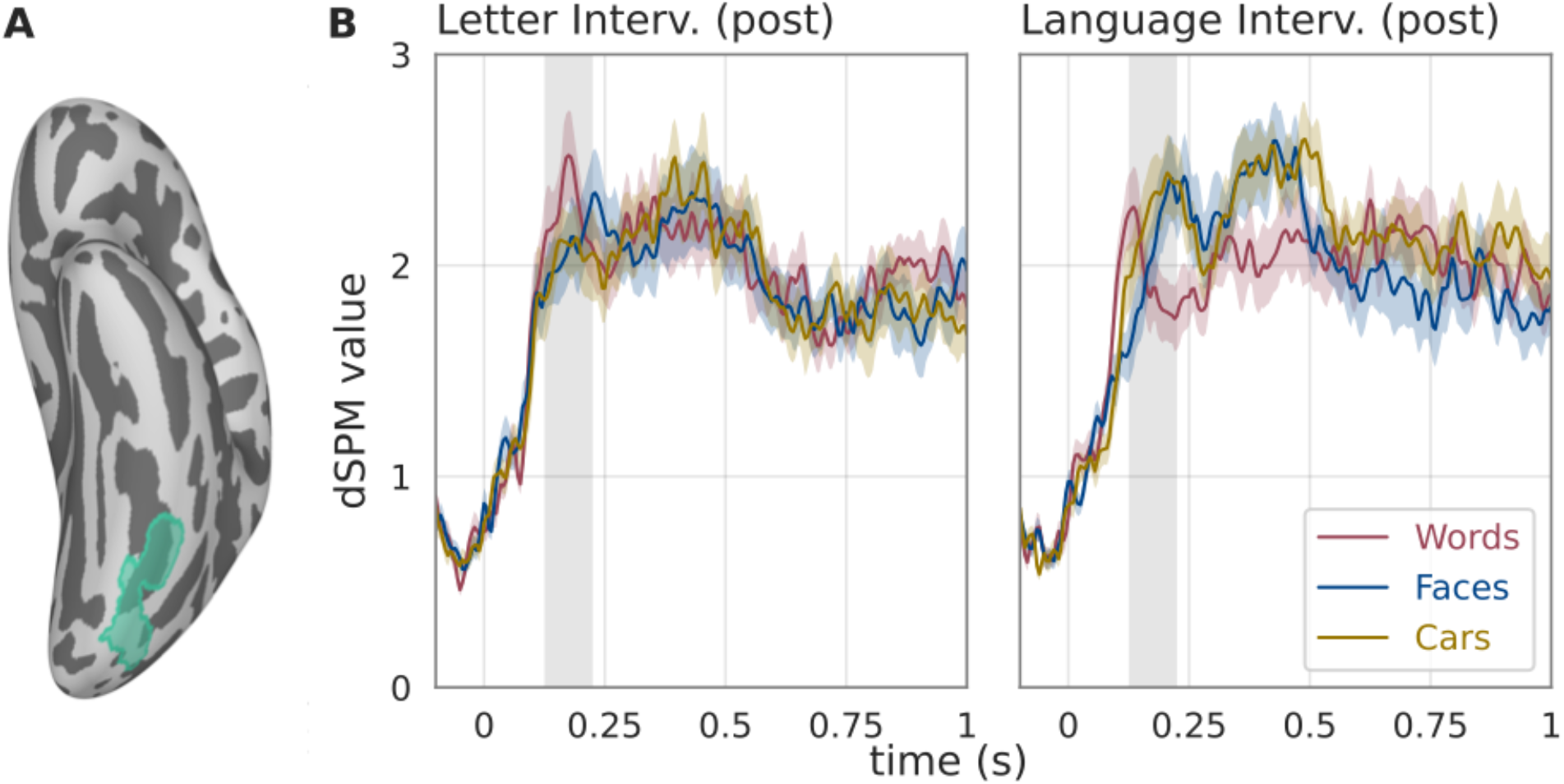
Word-selective responses differ between Letter and Language groups post-intervention. (A) Atlas-based region of interest (ROI) that selectively responds to words compared to other visual categories in literate adults (Rosenke et al., 2021). Source-localized MEG responses were extracted from this ROI and statistics were computed based on a 100ms time window defined around the peak of the grand average waveform (indicated with gray shading in panel B). (B) MEG responses to Words, Faces and Cars in the post-intervention dataset are shown for the two intervention groups. Only in the Letter Intervention group is there a word-selective response. In the Language Intervention group this patch of cortex shows a clear preference for images of faces and cars compared to words. Prior to the intervention, MEG responses did not differ between children assigned to the Letter and Language intervention groups.

To examine intervention-driven changes in tuning properties, we fit a LME model with fixed effects of group (Letter Intervention vs. Language Intervention), condition (Words, Faces, Cars dummy coded with Words as the reference condition), time (Pre-intervention vs. Post-intervention), and a full random effects structure (intercepts and slopes for all within subject effects (Barr et al., 2013)). The three way group, by condition, by time interaction revealed causal effects of the intervention on stimulus selectivity (F(2,126.7) = 4.67, p = 0.011). More precisely, the significant three way interaction indicates that the selectivity for words compared to other visual categories changes differently for children assigned to the Letter versus Language intervention program. The three way interaction reflected the change in the response to Words relative to Cars (t(126.7) = -2.97 p = 0.0036). There was not a significant three way interaction for Words compared to Faces (t(258) = -0.87, p = 0.38).

**Figure 3** shows responses to each stimulus category in the word-selective cortex ROI for the Letter and Language intervention groups and **Figure 4** shows how responses change in each intervention group. The pattern of intervention-driven changes provide some support for the hypothesis that the first phase of literacy learning involves competition between words and objects for cortical territory in VOTC (Kubota et al., 2019). Within a patch of cortex that broadly responds to images, the relative response amplitude to words relative to objects changed for children assigned to the Letter Intervention versus Language Intervention. However, examination of the pattern of changes in the Language Intervention group revealed unanticipated effects. A model comparing the change in MEG responses to words between the Letter and Language Intervention groups (group by time interaction) confirmed a significant two way interaction with the Letter group increasing and the Language group decreasing over the intervention period (t(126.7) = 2.43, p = 0.016).

### Literacy training rapidly establishes word-selective responses in visual cortex

We next analyzed pre-intervention and post-intervention data separately to investigate category selectivity before and after the intervention. Prior to the intervention, word-selective cortex was not selective for any stimulus category: A LME model of pre-intervention MEG responses with fixed effects of condition (Words, Faces, Cars), intervention group (Letter, Language), their interactions, revealed no main effect of condition (F(2, 88) = 0.43, p = 0.65). Moreover, as expected with random assignment, the two intervention groups did not differ: there was no main effect of group (F(1,44) = 0.96, p = 0.33) and no group by condition interaction in the pre-intervention data (F(2,88) = 1.27, p = 0.28). These results are in-line with previous reports that the patch of cortex that will eventually become the VWFA is broadly responsive to visual stimuli and not specifically tuned for a particular category in the pre-literate brain (Hervais-Adelman et al., 2019; Saygin et al., 2016).

**Figure 4:**
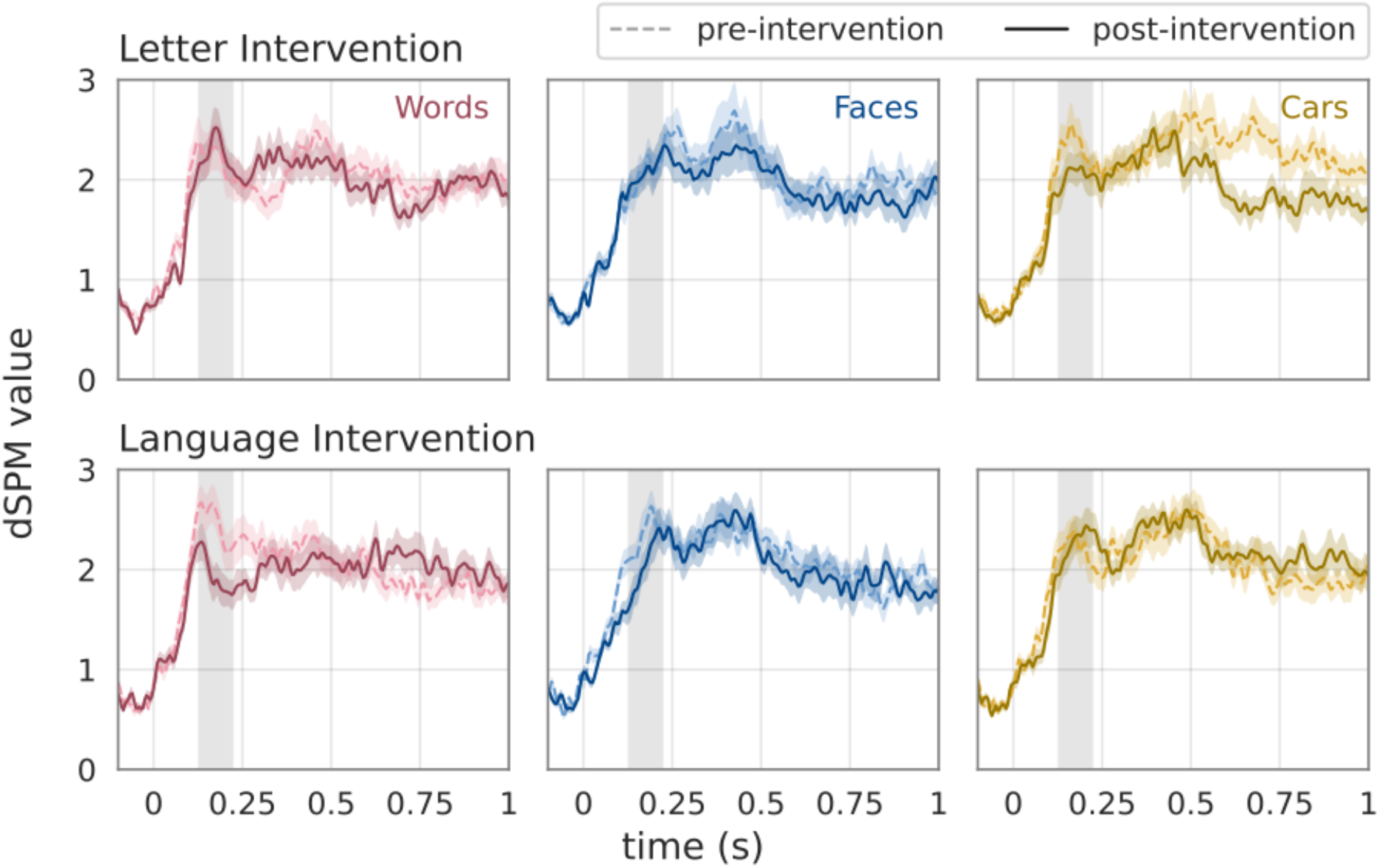
Changes in visual responses over the intervention. MEG responses to Words, Faces and Cars pre- and post-intervention for each group.

Fitting the same model to the post intervention data revealed a marginally significant group by condition interaction (F(2,88) = 2.83, p = 0.05 **Figure 3**). Compared to the Language Intervention group, the Letter Intervention group showed an enhanced response to Words relative to Cars (t(88) = -2.37, p = 0.02) but not Faces (t(129) = -1.34, p = 0.19). Thus, being assigned to the Letter versus the Language intervention caused differences in the tuning properties of category selective visual cortex.

### Do words compete with other visual categories for cortical territory?

To test the hypothesis that the Letter Intervention leads to an increased response to words at the expense of faces and/or objects we combined data from RCT described above with an independent replication cohort (n=16) that participated in the Letter Intervention (without random assignment). This replication cohort was originally conceptualized as a follow-up study, with a larger sample, specifically focusing on the neurobiology of reading acquisition. However, data collection was halted at the onset of the COVID-19 in March, 2020. Thus, the combined cohort included 40 participants who participated in the Letter Intervention and underwent the same MEG protocol. Individual MRI data was not available for all the participants in the replication cohort so the sensor data (n=40 participants combining original and replication cohorts) was aligned to a standard head position and statistics were calculated in sensor space.

We first examined changes in the evoked response for words by performing spatial-temporal clustering of the sensor-space data comparing the response to words post-versus pre-intervention. We found a significant increase in the response to words on left-lateralized posterior sensors spanning 300ms to 400ms after stimulus onset (**Figure 5 top panel;** p = 0.01). Source localization (based on coregistering each participant’s data directory to the fsaverage template) indicated that the effect was localized to VOTC (in the vicinity of the VWFA), lateral temporal cortex (superior temporal sulcus and middle temporal gyrus) and inferior parietal cortex (**Figure 5 middle panel**). The same analysis for faces and cars did not reveal any significant changes between the pre- and post-intervention data. Thus, any pruning of the response to faces and/or cars was not large enough and consistent enough to survive a spatial-temporal clustering correction for multiple comparisons.

**Figure 5:**
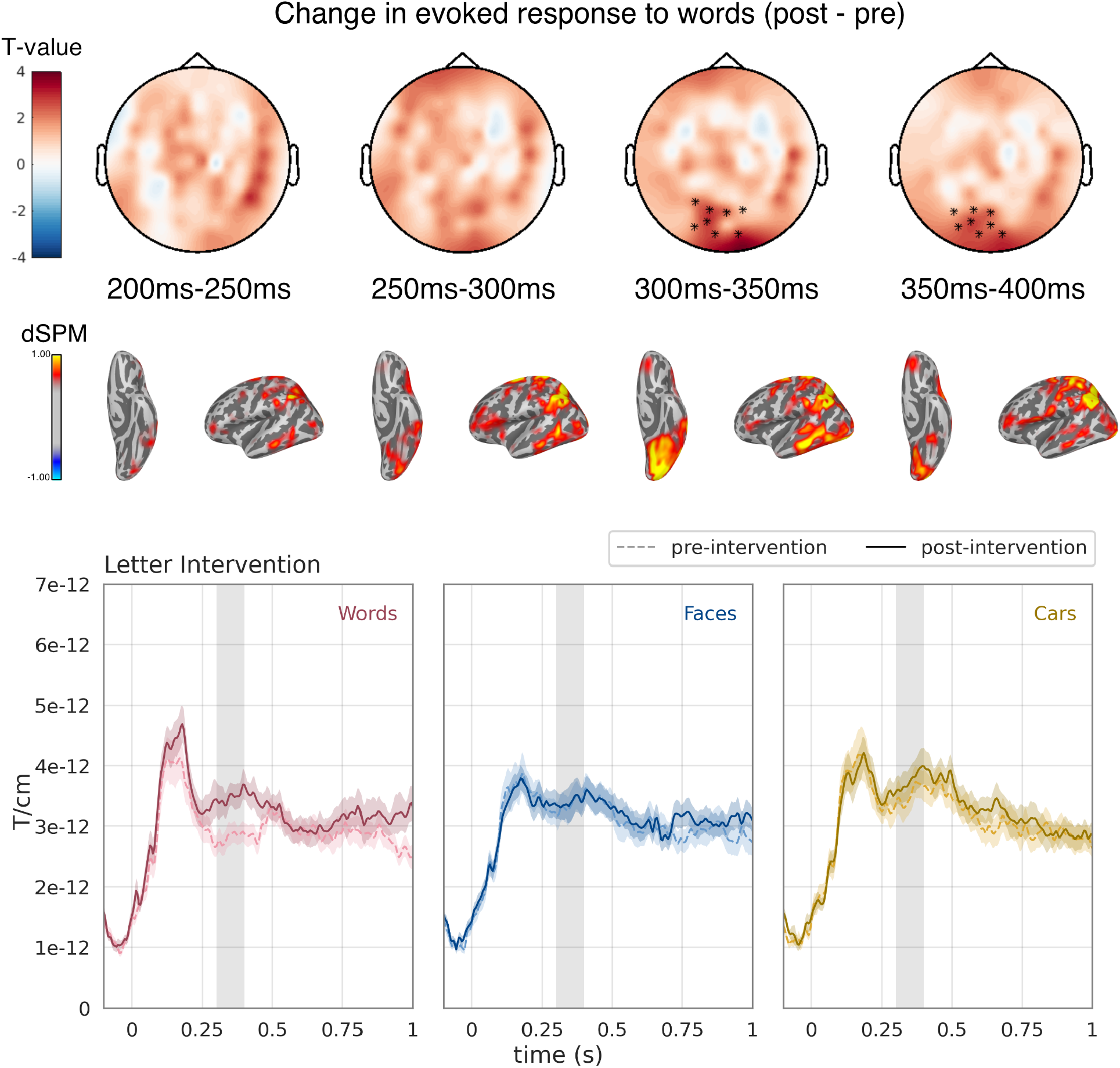
Enhanced response to words after the Letter Intervention. Topo plots show t-statistics comparing the response to words post-versus pre-intervention. Warm colors indicate an increase in response and asterisks indicate the cluster of sensors and timepoints that were significant based on spatiotemporal clustering of the sensor space data. Source estimates (dSPM values) are shown below with warm colors indicating an increased response to words after the letter intervention. The evoked response to Words, Faces and Cars is shown for the cluster of sensors with a significant intervention effect (gray shading indicates significant time window).

To more directly examine competition, we created a ROI encompassing the cluster of sensors showing a significant increase in the evoked response to words (shown in **Figure 5**). We then extracted the average evoked response within this ROI for each condition and time point and calculated the correlation between the change in response to words versus other categories. A negative correlation would indicate that the increase in word response was coupled to a decrease in the response to another category. We first examined the average response within the time window corresponding to the significant increase in the word response (300-400ms) and found a significant positive correlation for words and faces (r=+0.46, p=0.005) and a positive correlation that was not significant for words and cars (r=+0.27, p=0.11). We next conducted an exploratory analysis of 50ms time windows spanning 100ms to 400ms to ensure we hadn’t missed a time-localized negative correlation. For each time window there was a significant positive correlation between the intervention-driven change in the response to words and faces (+0.52< r <+0.65, 0.000015 < p <0.001). The correlation between the change in response to words and cars was positive at each time-point but was only marginally significant (+0.12< r <+0.37, 0.03< p <0.48). Thus, the increase in the response to words was not coupled to a decrease in the response to any other category within this ROI.

## Discussion

These findings represent the first causal test of neuronal recycling: pre-literate children were randomly assigned to two intervention programs, which were taught by the same teachers, employing the same didactic principles, in the same classroom, and with an analogous sequence of activities and play breaks, but focusing on two different skills, namely written language and spoken language. Response properties in high-level visual cortex changed over the two week intervention and being assigned to the Letter versus Language intervention produced different outcomes. Thus, visual cortex is much more plastic than often presumed: two weeks of basic literacy training begins prompting the emergence of word-selective responses. However evidence for competition between words and faces or objects for cortical territory was modest at best. In our analysis of the *a priori* VWFA ROI, we focused on an early time window corresponding to the bottom-up visual response and found that, for the Letter Intervention, there was an increased response to words relative to objects. This finding does provide some support for the notion that gaining expertise with new visual categories such as letters might recycle neuronal maps that would have otherwise been devoted to processing other categories of visual objects (Dehaene and Cohen, 2007; Kubota et al., 2019). However, other patterns of changes cannot be explained based on the neuronal recycling framework. For example, the Language Intervention group showed a decreased response to text (**Figure 4**). Moreover, a follow-up analysis with a larger cohort of Letter Intervention subjects did not reveal a significant decrease in the evoked response to faces or objects at any sensors or timepoints. In this larger sample there was a clear increase in the response to words but responses to other categories were stable (**Figure 5**). It is possible that the sensor space analysis lacked the sensitivity to detect the pruning of responses within a localized patch of VOTC. But it is also clear that the increased response to words was not coupled to a decreased response to other categories (see **Figure 5**). On the contrary, there was a positive correlation between changes in the response to words and faces indicating that, if anything, changes in the response to each category were linked as opposed to competing. This is in line with a cross-sectional study of adults which reported that literacy was correlated with an enhanced response to many visual categories without any indication of competition (Hervais-Adelman et al., 2019). However, a longitudinal study of school-aged children did observe words and limbs trading cortical territory (Nordt et al., 2021).

The present study focused on the initial phase of literacy learning where children begin learning to recognize letters, associate letters with sounds, and blend sounds to decode simple words. We focused on this initial phase of the learning process and employed a short but intensive intervention program to understand how early experiences in the classroom sculpt the neural circuitry that supports skilled reading for years to come. However, different mechanisms are most certainly at play at different stages of the learning process, and even though we didn’t observe strong evidence for competition over the timescale of weeks, these data do not rule out the possibility of competitive neural recycling over the timescale of years. Understanding the mechanisms at play over timescales ranging from hours to days, months and years is an important challenge for future work.

Based on our measures of intervention-driven changes in the functional architecture of visual cortex we can surmise that high-level visual cortex undergoes rapid changes as children enter school. How these changes play out over a longer timescale is still unknown but, in terms of neuronal recycling, there is no behavioral data suggesting that literacy has adverse consequences for any other aspects of visual perception (Hervais-Adelman et al., 2019; van Paridon et al., 2021). Instead, we might hypothesize that training the visual system to make fine-grained distinctions between complicated visual patterns leads to a modular structure that benefits perception more broadly. Indeed, the benefits of literacy on a myriad of brain functions ranging from basic visual perception, speech perception, language abilities and higher-level cognitive functions have been well documented (Dehaene et al., 2015, 2011; Huettig et al., 2018a, 2018b). The present study fills in a critical datapoint revealing changes in the tuning properties of category-selective visual cortex as young children are exposed to literacy.

## Materials and Methods

### Study Overview

48 English-speaking pre-school children (4y10m to 6y2m) were recruited for two-week language and literacy camps focusing on building foundational skills to prepare children for kindergarten. A parent or legal guardian provided informed consent and each child participant provided assent under a protocol that was approved by the Institutional Review Board at The University of Washington. All methods were carried out in accordance with the guidelines laid out in this protocol.

After enrolling in the summer program and providing consent/assent, participants were randomly assigned to one of two intervention groups: (1) The “Letter Intervention” which focused on letter identification and formation, grapheme-phoneme association, phonemic awareness, and blending three-letter, consonant-vowel-consonant (CVC) words; (2) The “Language Intervention” which focused on language skills such as listening comprehension, narrative structure, retelling oral narratives, syntax instruction, semantic feature analysis and vocabulary instruction. The Letter Intervention targeted the foundations of decoding skills following an approach grounded in systematic, multisensory, direct-instruction. The Language Intervention also employed systematic, multisensory, direct-instruction, however it targeted the compendium of language skills that form a critical foundation for reading development without any use of letters or other decoding related skills. The camps were held for 3 hours a day, five days a week and lasted two weeks totalling 30 hours of instruction in groups of six children and two teachers. Baseline behavioral testing, magnetic resonance imaging (MRI) and magnetoencephalography (MEG) sessions were conducted 19 days (SD = 9.23) prior to beginning the camp and follow-up sessions were conducted 8 days (SD = 3.90) after completing the camp for each child.

### Participants

All participants were native English speakers who were planning to start kindergarten in the fall of 2019. 47 out of the 48 participants had attended preschool/pre-k, public, or private daycare. Participants from the Letter (N=24) and Language (N=24) intervention both participated in the same intake session with the same recruitment criteria. No diagnoses of ADHD, ASD or any other developmental disorder was reported. Everyone demonstrated normal or corrected-to-normal vision. Participants were not aware that there were two different interventions.

### Sample size and statistical power

Since the present study is the first randomized controlled trial (RCT) of learning-induced changes in the brain’s response to text, we did not have any equivalent designs to use for a power analysis. Thus, a power analysis was conducted based on previous work that related EEG measures of text-selective responses to letter knowledge in preschool children of the same age range (Lochy et al., 2016). Lochy and colleagues used a between group design to compare the amplitude of the text-selective response in preschool children with high, versus low, alphabet knowledge. They reported the mean amplitude for children with high alphabet knowledge as 1.770 µV (SD=0.262), versus 0.657 µV (SD=0.214) for children with low alphabet knowledge. This corresponds to an effect size of Cohen’s d=4.65 meaning that even for a sample of 20 subjects, the statistical power is above 0.95. Given the large effects sizes in cross sectional data, we proposed a sample size of N=44 (n=22 per group) in the clinicaltrials.gov registration (NCT03945097) and we exceeded this target with a final sample of N=48 (n=24 per group).

### Overview of the Language and Literacy Camp Intervention Programs

We designed the “Language and Literacy Camp” intervention programs to train children on foundational academic skills over the course of short (2 week), engaging summer camps. Intervention activities were provided by 3 teachers with a master’s or bachelor’s degree in Education and/or Speech Pathology, and with prior experience teaching young children. Before beginning intervention activities, two pilot camps were run, each four days with 4 children. These pilot camps were used to fine tune logistics and curriculum details. The pilot camps also provided practice, training and time for coordination between the three teachers to ensure consistent practices throughout the duration of the program.

Interventions took place in small groups (6 children and 2 teachers per camp) in a preschool classroom. Interventions occurred in the morning (9am-12pm) or afternoon (1pm-4pm). A minimum attendance of 8/10 days was required for each participant. Most participants attended all days, and everyone attended at least 8 days. Before enrolling, parents agreed to bring their child to each session barring illness or unavoidable events. Both camps were designed similar to a traditional half day kindergarten classroom with the following schedule (every day of class, except for the first day, followed the same structure; Table 1).

**Table 1:** Daily schedule for the Letter and Language Intervention programs. *Replaced by introduction and rule setting on day 1.

| Time | Letter intervention | Language intervention |
| --- | --- | --- |
| 9:00 AM / 1:00 PM | Welcome + games | Welcome + games |
| 9:20 AM / 1:20 PM | *Phonological awareness \** | *Morphology and Syntax\** |
| 9:45 AM / 1:45 PM | Recess | Recess |
| 10:00 AM / 2:00 PM | *Direct Instruction part 1* | *Oral Story Retelling* |
| 10:25 AM / 2:25 PM | Centers | Centers |
| 10:45 AM / 2:45 PM | Recess/snack | Recess/snack |
| 11:05 AM / 3:05 PM | *Direct Instruction part 2* | *Vocabulary* |
| 11:30 AM / 3:30 PM | Centers | Centers |
| 11:50 AM / 3:50 PM | Story | Story |
| 12:00 PM / 4:00 PM | End of class | End of class |

### Magnetoencephalography

Magnetoencephalography (MEG) and magnetic resonance imaging (MRI) data was collected for each participant immediately before and after the intervention program. The MEG experiment was designed to probe the tuning of VOTC to different categories of visual images and MRI data was used for individual source reconstruction.

#### Stimuli and experimental procedure

All the procedures were controlled by in-house Python software (expyfun: https://github.com/LABSN/expyfun) and stimuli were displayed on a gray background (50 cd/m2) of a back-projected screen using a PT-D7700U-K (Panasonic) projector. All code to reproduce the experiment can be found at: https://github.com/YeatmanLab/SSWEF/. **Figure 1** shows the procedure of the experiment. Children were introduced to the game “Alien Adventures” where they had to search for the hiding alien (oddball). Children were instructed to maintain fixation at the center of the screen and press a button each time they saw the friendly alien. The experimental stimuli consisted of grayscale images of: a) pronounceable pseudowords rendered in an unfamiliar font (“Words”); b) childrens’ faces, and c) cars. All images were rendered on a textured background to control for differences in visual field coverage and the images were manipulated with the Shine toolbox (Willenbockel et al., 2010) such that images from each category were matched in terms of (1) contrast, (2) luminance, and spatial frequency power distribution (Stigliani et al., 2015). Importantly, the text was rendered in an unfamiliar manner with textures filling the letters, and the shape of the word stretched at various angles. This ensured that the participants did not have any specific familiarity with the word images. The faces and cars were also at variable locations and angles within the image. Cars were chosen as the category of objects for three reasons: a) children have familiarity with cars but, at this age, do not have specific expertise with cars; b) images of cars produce robust activation throughout object-selective regions but they do not have a devoted region for the subordinate category; c) our previous work suggested that cars make a good comparison condition for words to index the development of word-selective responses (Kubota et al., 2019). Thirty images were randomly drawn from each category and, on each trial, the image was displayed for 1.0s followed by a blank screen with a random duration between 0.5s and 1.0s sampled from a uniform distribution. In addition ten oddball images (colorful aliens) were inserted at random points in the experiment and these trials were removed from the analysis. Children performed well on the target detection task (mean d’ = 3.3, range 1.2-4.2) confirming that they were attentive to the stimuli.

#### MEG and MRI data acquisition

MEG data were recorded inside a magnetically shielded room (IMEDCO) using a 306-channel dc-SQUID VectorView system (Elekta-Neuromag). Neuromagnetic data were sampled at 1kHz with a passband of 0.01 to 600 Hz. A 3D position monitoring system (Polhemus, Colchester, VT) was used to record the locations of head position indicator (HPI) coils, cardinal (nasion, left/right preauricular) anatomical landmarks, and at least 100 digitized scalp points (which were used to coregister the MEG sensors with individual structural MRI). HPI coils were used to record the subject’s head position continuously relative to the MEG sensors, to allow offline correction of movement-related artifacts.

Individual structural MRIs were obtained at The University of Washington Diagnostic Imaging Science Center (DISC) on a Philips Achieva 3T scanner using an 8-channel phased-array SENSE head coil. A whole-brain anatomical volume at 0.8 × 0.8 × 0.8 mm resolution was acquired using a T1-weighted MPRAGE sequence (TR 15.22 s, TE 3 ms, matrix size 320 × 320, field of view 240 × 256 × 169.6, 212 slices). Head motion was minimized by an inflated cap, and participants were monitored through a closed-circuit camera system.

#### MEG data preprocessing, source reconstruction and statistics

MEG data were analyzed using MNE-Python (Gramfort et al., 2013). Code to reproduce all analyses can be found at: https://github.com/YeatmanLab/SSWEF/. Environmental noise reduction was performed using MNE-Python’s Maxwell filter function and data were individually corrected for head movements using the average of each participant’s head positions as a reference (Taulu and Kajola, 2005; Uutela et al., 2001). The temporally extended signal space separation method was applied with a correlation limit of 0.98 and a segment length of 20 seconds (Taulu and Hari, 2009; Taulu and Simola, 2006). Bad channels were substituted with interpolated values. Next the neuromagnetic data were low-pass-filtered (40 Hz cutoff), and signal space projection was used to suppress cardiac artifacts identified using peripheral physiologic (ECG) sensor data and ocular artifacts using EOG sensors (Uusitalo and Ilmoniemi, 1997).

To determine trial rejection thresholds, we developed a data driven approach that maximizes the reliability of the measurements (**Figure 6**). Briefly, this involved a grid search over a wide range of peak-to-peak amplitude rejection thresholds for magnetometer and gradiometer signals, designed to maximize the mean correlation across subjects between pre- and post-intervention evoked responses in bilateral early visual areas.

**Figure 6:**
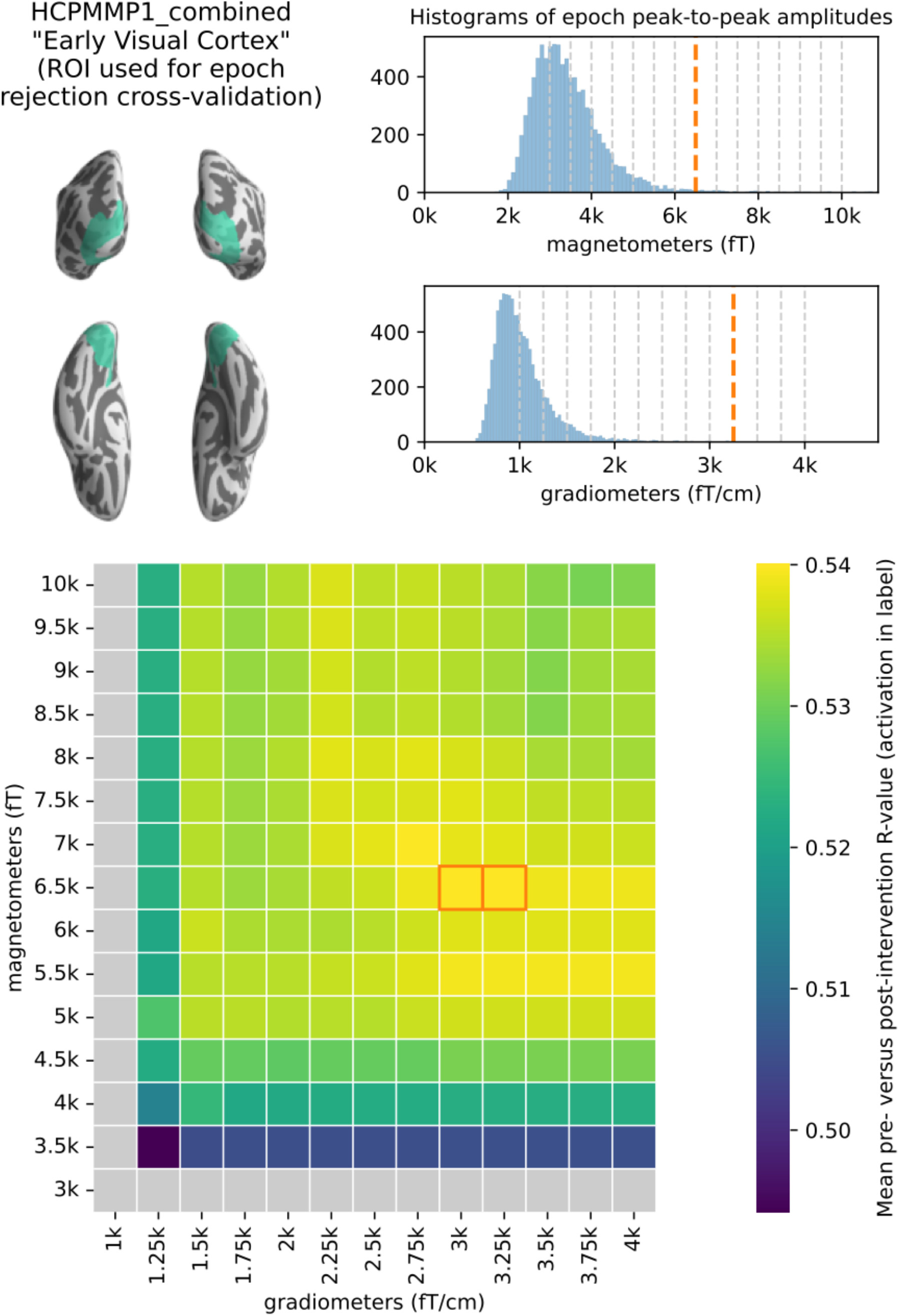
Procedure for determining epoch rejection thresholds. (A) Early visual cortex region of interest (ROI) from which average evoked signal was extracted from pre- and post-intervention grand averages (over trials) for each subject. This ROI was selected for tuning epoch rejection thresholds in order to maximize the reliability of the evoked response in early visual cortex. (B) Histograms of pre-rejection peak-to-peak amplitudes across all subjects and trials. Pale dashed vertical lines indicate grid search locations, thick and dark dashed lines indicate the selected rejection thresholds. (C) Heatmap of grid search results; each cell is the mean (across subjects) correlation between pre- and post-intervention evoked activity in the ROI shown in (A). The cells corresponding to the optimal thresholds are outlined; the higher (more permissive) of the two was chosen. Cells with pale gray color indicate untenable thresholds (those resulting in rejection of all trials in at least one experimental condition for at least one subject). Through this procedure we determined epoch rejection thresholds that led to a reliable evoked waveform in early visual cortex.

The resulting MEG signal data were windowed into 1100ms epochs of evoked neuromagnetic activity (including 100ms pre-stimulus baseline.) Evoked trial data were DC drift corrected using a mean baseline correction approach. Event related fields (ERFs) were obtained by averaging the remaining artifact-free trials of each participant and condition. Data from 3 participants was discarded due to technical problems with cHPI or MEG recordings.

For individual subject MEG source reconstruction, each participant’s T1-weighted MPRAGE image (selecting the highest quality of the participants T1s) was segmented with Freesurfer (Dale et al., 1999; Fischl, 2012). Then an anatomically constrained three-compartment boundary element model (BEM) consisting of the inner skull, outer skull, and scalp surfaces was used as a conductor model. A source space of current dipoles was constructed from the high-resolution tessellated pial cortical surface using a recursively subdivided icosahedron, yielding 10242 dipoles per hemisphere.

Dipole source orientations were unconstrained to account for eventual surface warping between child cortical surfaces and a template brain based on adult subjects (see below). The resulting individualized BEM and source space was used to provide an accurate forward solution mapping dipole currents in the source space to the recorded ERFs. Using this colocation information and an iterative L2 minimum-norm linear estimator with shrunk noise covariance from the baseline sensor covariance (Engemann & Gramfort, 2015) we computed dynamic statistical parametric maps (dSPM) of conditional ERFs (Dale et al., 2000). For group level analysis individual dSPMs were mapped to the FreeSurfer fsaverage cortical template using a non-linear spherical morphing procedure (20 smoothing steps) that optimally aligns individual sulcal–gyral patterns (Fischl et al., 1999).

To limit the number of statistical comparisons in space and time, we focused on an *a priori* region of interest (ROI) defined on the fsaverage template. We used the maximum probability map of word-selective visual cortex from (Rosenke et al., 2021). Specifically, we combined the posterior occipitotemporal sulcus (pOTS), inferior occipital sulcus (IOS) and inferior occipital gyrus (IOG) ROIs into a single region. We then aggregated source-localized responses (dSPM values) within this region using the first right-singular vector of a singular-value decomposition of the sources, scaled to match the mean per-vertex power and sign-flipped to match the dominant direction of sources in the ROI, resulting in a single time course for each ROI. Since our main interest was in the visual evoked response to the different image categories, we then identified the first peak in the grand average waveform for this ROI (which occurred at 175ms) and then defined a 100ms window around the peak (125ms to 225ms). All statistics were computed on the average evoked response within this time window for the pOTS/IOS/IOG ROI. All statistical analyses were conducted using linear mixed effects models with a maximal random effects structure (all within-participant effects were included as random effects in line with the guidelines laid out in (Barr et al., 2013; Bates et al., 2007)).

### Behavioral testing

Each participant underwent three behavioral testing sessions: an intake session to assess eligibility for the study, pre-intervention and post-intervention behavioral assessments focusing on reading and language skills.

#### Intake session

During the intake screening session the following tests were administered:

1. Peabody Picture Vocabulary Test (PPVT) 4th Edition (Dunn and Dunn, 2007).
2. Uppercase letter knowledge: Flash cards for 26 upper case letters. Each card was shown to the child and they were asked “What letter is this?” and “What sound does it make?”.
3. Preschool Word and Print awareness (Justice and Ezell, 2001).
4. Comprehensive Test of Phonological Processing Rapid Automatized Naming: Colors and Shapes (Wagner et al., 1999).
5. Test of Preschool Early Literacy (TOPEL) – Subtest 3: Phonological Awareness (Lonigan et al., 2007).
6. Edinburgh Handedness Questionnaire (Oldfield, 1971).

After the testing was complete, each child was taken to a replica MRI scanner to practice holding still during scanning. Finally, children completed the Snellen Eye Chart to assess vision. Children were invited to participate based on the following criteria:

1. Did not know all uppercase letters and sounds
2. PPVT standard score ≥ 86
3. Normal or corrected to normal vision
4. Was comfortable in the mock MRI and able to hold still for 5 minutes

#### Pre- and Post-Intervention Sessions

The following tests were always administered in the same order:

1. Letter knowledge uppercase: Flash cards were used to assess knowledge of all uppercase letters of the alphabet with their accompanied sound in a random order.
2. Phonological and Print Awareness Scale (PPA): Initial sound matching, Final sound matching, Phonemic awareness (Williams, 2014).
3. Letter knowledge lowercase: Flash cards were used to assess knowledge of all lowercase letters of the alphabet with their accompanied sound in a random order.
4. Phonological Awareness Literacy Screening (PALS): Pseudoword decoding list (Invernizzi et al., 2003).
5. Narrative Language Measures: Story Retell (Petersen and Spencer, 2012).
6. Expressive Vocabulary Test Third Edition (EVT).

### Letter Camp Intervention Curriculum

The Letter Intervention implemented a sequence of well-established early reading instructional activities (Castles et al., 2018) in a two-week curriculum, based on pedagogical models of Direct Instruction and Gradual Release of Responsibility (Fisher and Frey, 2013). Intervention activities consisted of phonological awareness, letter identification, letter production, phoneme grapheme association, and CVC non-word blending/reading (Table 2). Children learned two letters a day. For each letter, students gradually took more responsibility for their own learning, starting with teacher-led, direct instruction and transitioning to more independent activities on their own or in pairs. Each lesson began with a review of the material learned from previous days so that learning was scaffolded by a strong foundation.

The curricula were developed by a speech-language-pathologist and a teacher with many years of teaching experience in typically and atypically developing children of this age group. All materials were developed in the lab or obtained via free websites (e.g. http://www.floridaearlylearning.com). Images were obtained from free clipart databases such as http://clipart-library.com and https://thenounproject.com. The direct instruction of the Letter camp was partially based on the Slingerland Approach (Reading, n.d.), a classroom adaptation of the Orton Gillingham method (Orton, 1966; Ritchey and Goeke, 2006). The teacher leading direct instruction was officially trained in the Slingerland method.

#### Phonological awareness (PA)

After 20 minutes of free play at the start of each day (blocks, puzzles, modeling clay), whole-group phonological awareness instruction took place. Phonological awareness activities included songs, games, and gross motor movement. Lessons progressed in difficulty, beginning with larger phonological units and continuing to smaller phonological units over the course of ten days: syllable segmentation, syllable blending, onset-rime blending, onset-rime segmentation, C-VC blending, C-VC segmentation, CVC blending, and CVC segmentation. Each lesson began with a review of the previous lessons to reinforce the previously taught skills.

#### Direct Instruction in Letters

Two lowercase letters were taught each day (one letter at a time) using direct instruction. Following the Slingerland method, the teacher first introduced the name and shape of the target letter using a letter-picture association card. The teacher modeled how to form the letter on a whiteboard (including verbal directions about where to start forming the letter, stroke direction, etc). After modeling, each student formed the letter on the whiteboard, with pre- and post-reinforcement of the letter name (“What letter are you going to write?” / “What letter did you write?”), and verbal directions during the activity. The teacher scaffolded the student by guiding proper pencil grip, controlled motor movement, stroke direction, etc. Positive feedback in the form of praise and encouragement as well as recognition of effort was provided to each student (Mangels et al., 2006).

Following the whiteboard activity, students used printed letterform diagrams (a dot at the starting point, arrows showing stroke direction, etc) to trace the letter shape with their fingers. While tracing, students verbalized the name of the letter while forming it on paper. If necessary, teachers provided individual guidance on pace, start point, directionality, smooth motor control and integration of the student’s hand, arm and shoulder. Following finger tracing, students traced the letter using the eraser side of a pencil, and finally with the sharpened end of a pencil, again verbalizing the letter name while tracing its shape. At each phase of the activity, students received feedback on directionality, pace, pencil grip, starting point, integration of hand, arm, and shoulder movements, and smooth motor control (as needed). One of the goals of this method is attuning students to a multisensory representation - visual form, kinesthetics and sound - helping them remember all the aspects of the letter.

Following letter formation, students returned to the group to begin grapheme-phoneme association. Again, following the Slingerland approach, the teacher showed the letter picture association card and stated “This is the letter. It makes the sound you hear at the beginning of this picture.” The teacher asked the students to name the picture and the teachers and students worked to hear the first sound of the picture. After identifying the first sound of the picture, the students practiced saying the sound and the teacher also asked each student to individually produce the sound while checking for articulation and proper pronunciation. Finally, the teacher instructed “We talk about the letter like this: “<a> apple [æ]” while writing the letter in the air at the same time. The class practiced “talking about” the letter as a group and then each child practiced individually. Finally, the students reviewed letter flashcards, while talking about all previous learned letters, including the current one.

#### Direct Instruction in Blending and Decoding

Starting at day 3, when sufficient letters were learned to form words, CVC sequence decoding was included after the daily letter review. To explicitly teach decoding the teacher modeled how to identify the first letter name and sound, then the second letter name and sound, and then the third. With this information, the teacher modeled how to blend the 3 sounds together to read the CVC sequence. Each student was given a chance to decode at least two words by themselves. Depending on the level of support each student needed, the teachers scaffolded the process so that the student could successfully blend sounds and decode the word. After decoding the sequence, the student had to decide if it was a real or a nonsense word. Each student was applauded with praise, encouragement, enthusiasm and recognition of effort (Yeager et al., 2019).

#### Center activities encompassing phonological awareness, letter and reading instruction

Learning stations were set up around the classroom to give students hands-on experience with the new letter and sound material they had learned. Each station reinforced prior learning and included high interest material such as letter matching games, dot markers, duplo legos, fishing rods, stuffed animals, crayons, mystery sound boxes, blocks, modeling clay and other developmentally appropriate activities. Students worked individually or in pairs at 4 stations per session, spending 6 minutes at each station. During these rotations, each student rotated to at least one teacher-led station for individual instruction on phonological awareness or CVC word blending.

#### Story time

At the end of the day a brief, 5-10 minute, story was read by the teachers. The story (content or reading comprehension aspects) was not further discussed, in contrast to the Language camp stories.

### Language Camp Intervention Curriculum

Similar to the Letter Camp Intervention, the Language Camp Intervention also implemented direct instruction and gradual release of responsibility pedagogical models for all activities. It included instruction in English syntax, narrative text structure analysis, story retelling, and vocabulary. All activities incorporated oral language and listening comprehension skills; no letters, phonological awareness or decoding were present in any language camp activity.

#### Color-coded grammar and Syntax Awareness

After 20 minutes of free play at the start of each day (blocks, puzzles, playdough), whole group syntax instruction occurred for 20 minutes. The instruction was designed to be multisensory, providing combinations of auditory, visual (colors and images) and kinesthetic (touching/moving of cards) interactions (Balthazar et al., 2020). This multimodal approach serves an effective learning environment (Shams and Seitz, 2008), with a high level of participation and enjoyment for the children, and opportunities to engage with the material through a variety of modalities. The participants were introduced to syntax by the function of words, without explicitly naming them (e.g. the child learns what an adjective is and how to use it, but will refer to it as “yellow cards” instead of using the term “adjective”; **Figure 7**). This color-coding approach has been proven effective in classroom programs (Goossens et al., 1992), second language learning (Kohler, 2009), for children with specific language impairments (Balthazar et al., 2020) and even adults with impaired language functions (Ebbels et al., 2014; Newton et al., 2017).

**Figure 7:**
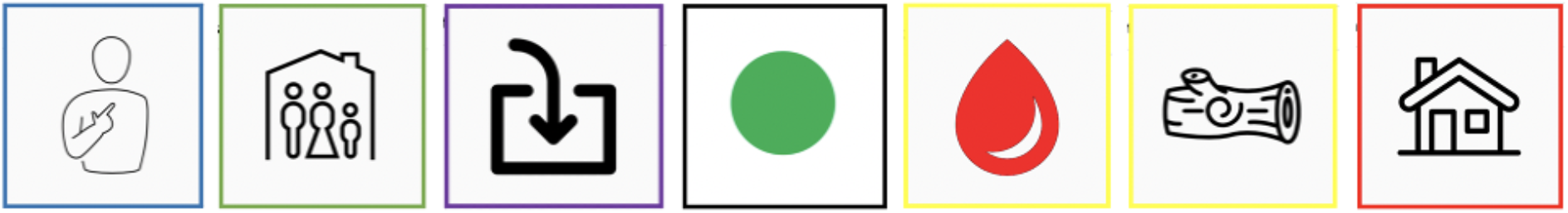
Example of syntax exercise with color-coded cards representing the different functions of words. The sentence represented here is “I live in a red wooden house”. This activity was introduced on day 3, when children learned about adjectives. Words that were hard to visualize like e.g. “a” were given a symbol that stayed identical throughout the camp. For example, I (personal pronoun image card) - live (verb image card) - in (preposition image card) - a (a/the symbol card) - red (adjective image card) - wooden (adjective image card) - house (noun image card)).

Syntax instruction gradually increased in complexity each day. It started with the idea of what a phrase is, and later introduced parts of speech: nouns, verbs, adjectives, subject verb agreement, verb tense, possessives, articles, prepositions. The goal was for children to learn how to effectively use grammar in sentences and color-coding was used to scaffold learning (Kohler, 2009). One card, with a distinctive color and image were used per word to build the sentences. Every card with a distinct word function (e.g noun, verb, adjective) had a different color. Images were used to visualize words.

Before each syntax lesson, the teacher reviewed cards from prior lessons for reinforcement and practice. Each day, the teacher introduced a new syntax concept. The vocabulary was introduced at the beginning of every session and cards were reused for multiple days to be able to focus on syntax over vocabulary. The teacher introduced the new syntax by incorporating the new (and old) cards into a sentence structure. As a whole group, students practiced verbalizing the syntax structure using the cards set out by the teacher. Lastly, each student took a turn producing a syntax structure using different cards.

#### Narrative comprehension

Understanding and retelling stories lays a foundation for comprehending more complex narratives throughout life (Calfee and Patrick, 1995). During a 25-minute lesson, students received explicit, systematic instruction and guided practice in the following story elements throughout the first week: characters, setting, problem and solution, lesson/theme, order of a story (beginning, middle, end) (Archer and Hughes, 2011). Using age-appropriate anchor texts (e.g., *The Three Little Pigs*) (Calfee and Patrick, 1995), students listened to a story read by a teacher, then discussed the story elements they learned. The teacher introduced the story element, showed the students where it occurred in the story and students identified the same element, based on structural questions (Morrow, 1984). Before each new lesson, the teacher reviewed previously taught story elements. In the second week of the camp, variations of anchor stories were used to allow the children to further practice identifying all story elements learned in the first week.

#### Story vocabulary

Vocabulary was taught through contextual vocabulary instruction in which word context and personal experience are used, in combination with other information about the word, to teach the meaning of the complex words to children (Rapaport, 2005). A complex (“tier 2”), vocabulary word in the daily anchor story was introduced to the children, based on The Florida Center for Reading Research (*FCRR) New Vocabulary Instructional routine (e*.*g;* https://fcrr.org/sites/g/files/upcbnu2836/files/media/projects/empowering-teachers/routines/pdf/instRoutines_KV.pdf). Children were introduced to the word in a sentence, learned the meaning of the word, and learned to use it in a new sentence themselves. Previous words and their meaning were rehearsed daily.

#### Vocabulary training (themes)

The theme-based vocabulary training was grounded in the idea that providing children with ‘root words’ will establish a foundation for language learning. Four research-based vocabulary teaching practices were implemented (Christ and Wang, 2010): (1) Provide purposeful exposure to new words, (2) Intentionally teach word meanings, (3) Teach word-learning strategies, (4) Offer opportunities to use newly learned words that help young children acquire new vocabulary.

A direct instruction approach was implemented to learn vocabulary by theme. Throughout the two weeks, words under the following categories were taught: shapes and colors, body parts and positions, clothes, animals, food, transportation, sports and careers. Days 5 and 10 served as opportunities to review previously taught vocabulary. The vocabulary words were age appropriate, and included words just above and below the children’s age level. Including vocabulary at varying levels gave children the opportunity to acquire easy words and motivate them to learn more difficult words. Every day four different exercises were conducted during direct instruction using picture cards for each word: 1) The vocabulary cards were spread out and the students guessed the topic of the day. Subsequently students named the cards (fulfilling teaching practice 1 and 2 noted above). 2) The plural form of the vocabulary was practiced. The standard sentence *“I have one …, but I want two …*.*”* was continuously used throughout the program (fulfilling teaching practice 3 and 4 noted above). 3) The children had to find categories and networks within the theme (fulfilling teaching practice 1 through 4). This could be directly related to previously learned vocabulary (e.g. categorize clothes by body parts) or introducing some new, extra vocabulary. 4) The final step was a combination exercise that varied every day (fulfilling teaching practice 4). To finish up the day, the children repeated all the new vocabulary of that day out loud.

#### Center activities encompassing narrative comprehension and vocabulary training

##### Round 1: Narrative comprehension

The first center rotation each day reinforced prior learning and included high interest material such as puppet theaters with puppets from anchor texts, books on tape, pretend play with props from the anchor texts (characters, setting etc.), retelling practice, sequencing picture cards from the anchor text from beginning to end, and realistic fiction picture card scenes from beginning, middle to end, giving children hands-on experience with the new material they had learned. In week two, the main goal of the center activities was to get the children to retell a complete story. During center rotations, groups of two were formed, so children would be able to learn from each other and help each other. Story retell is a new skill for many children of this age and requires scaffolded practice. Each center activity lasted 6 minutes each with a total of 4 rotations per session.

##### Round 2: Vocabulary training

The second center rotation each day reinforced practicing and automatizing the introduced vocabulary. Center activities involved picture sorting, categorizing, drawing, or playing a game with new words (memory, bingo, puzzles, worksheets). During center rotations, each student rotated to at least one teacher-led center for individual practice on the vocabulary theme. Each center activity lasted 6 minutes each with a total of 4 rotations per session.

#### Story time

Story time in the Language Intervention Camp was different from story time in the Letter Intervention Camp. Participants were asked to apply the learned aspects from the narrative comprehension curriculum to the specific story. For example, on day 1 the story element “characters” was introduced. After story time the students were asked to identify all characters in the story. This expanded every day by adding new story elements.

### Replication Cohort

In addition to the RCT, additional data was collected on an independent replication cohort that participated in the Letter Intervention (without random assignment). This replication cohort was originally recruited with the goals of a) replicating the results from pre-registered RCT and b) including a larger sample size to investigate the neurobiological underpinnings of learning differences. However, data collection was halted at the onset of the COVID-19 in March, 2020 after 16 participants had completed the Letter Intervention. The replication cohort participated in the same MEG protocol but individual MRI data was not available for all the participants in the replication cohort so data was analyzed in sensor space.

MEG data from the 16 participants of the replication cohort was combined with data on the 24 RCT participants who were assigned to the Letter Intervention for a combined sample of 40 participants. Sensor data was aligned to a standard head position and statistics were calculated in sensor space. Rather than focusing on an *a priori* ROI and time window, we capitalized on the larger sample to examine changes in the response to words, faces and objects across all sensors and timepoints between 100 and 400 ms using the spatiotemporal clustering algorithm (Maris and Oostenveld, 2007).

## Acknowledgements

We would like to thank the families that participated in this study along with the funding sources for this work: NICHD R01HD095861, R21HD092771 and the Bezos Family Foundation.

## Author Contributions

JDY conceptualized the study and wrote the manuscript; JDY, LG and SE designed the intervention programs; SE, LG and MET ran the intervention programs; JDY and SJJ designed the MEG experiment; JDY, DRM, SC, ERP, EL, ST, and MC analyzed the data; JDY, ERP, GOB, ECK, LG, SE, and MET collected the data; PKK, ST, and EL provided guidance throughout the study.

## Data and Materials Availability

All data and code used for this paper will be made publicly available upon publication.

## Supplementary Information

**Supplementary Table 1:** Detailed Schedule for a day of Letter Intervention

| Time | Activity | Example |
| --- | --- | --- |
| 15 minutes | Free Play | puzzles, blocks, playdough |
| 25 minutes | Phonemic Awareness | Song, games, chant, etc. |
| 25 minutes | Recess | Free play outside |
| 4 minutes | Direct instruction part 1 (new letter) | Teacher introduces letter picture card.<br>Teacher models letter formation on white board.<br>Students individually trace letter at white board. |
| 3 minutes | letter formation practice | Students trace letter on 11X17 paper using 2 fingers, eraser, and pencil tip. |
| 3 minutes | phoneme grapheme correspondence | Teacher introduces key word and letter sound.<br>Teacher and students air write letter and say its "name, key word and sound"<br>Class reviews letter flashcards |
| 5 minutes | decoding | Students decode a cvc word at the white board with teacher assistance |
| 25 min | center rotation | games, coloring, tracing, sorting, matching, etc. |
| 25 minute | snack and recess | free play outside |
| 4 minutes | Direct instruction part 2 (new letter) | Teacher introduces letter picture card.<br>Teacher models letter formation on white board.<br>Students individually trace letter at white board. |
| 3 minutes | letter formation practice | Students trace letter on 11X 17 paper using 2 fingers, eraser, and pencil tip. |
| 3 minutes | phoneme grapheme correspondence | Teacher introduces key word and letter sound.<br>Teacher and students air write letter and say its "name, key word and sound"<br>Class reviews letter flashcards |
| 5 minutes | decoding | Students decode a cvc word at the white board with teacher assistance |
| 25 min | center rotation | games, coloring, tracing, sorting, matching, etc. |
| 10 minutes | read aloud | Teacher reads story to class |

**Supplementary Table 2:** Detailed Schedule for a day of Language Intervention

| Time | Activity | Example |
| --- | --- | --- |
| 15 minutes | Free Play | puzzles, blocks, playdough |
| 25 minutes | Syntax Instruction | Noun, verb, plural, tense |
| 25 minutes | Recess | free play outside |
| 5 minutes | Read aloud | Narrative Text |
| 10 minutes | Narrative Text<br>Instruction | Character, setting, etc. |
| 25 minutes | Center Rotation | games, coloring, tracing, sorting, matching, etc. |
| 25 minutes | Snack and Recess | free play outside |
| 15 minutes | Semantic Feature<br>Analysis Instruction | Colors, shapes, jobs, clothing, etc. |
| 25 min | Center rotation | Games, coloring, tracing, sorting, matching, etc. |
| 10 minutes | read aloud | Teacher reads story to class |

